# Semantic elaboration determines the time course of alpha-beta oscillations during the encoding and retrieval of narrative memories

**DOI:** 10.1101/2025.08.21.671460

**Authors:** Emmanuel Biau, Francesca M. Branzi

**Affiliations:** Department of Psychology, Institute of Population Health, University of Liverpool, UK

**Keywords:** Semantic processing, Episodic memory, Alpha-beta oscillations, Electroencephalography (EEG)

## Abstract

Understanding and remembering everyday events requires us to retrieve semantic relationships between elements by activating prior knowledge, allowing us to form a deeper memory trace. However, the neural interplay between semantic and episodic systems to encode and retrieve recent memories remains unclear. The present study addresses how semantic elaboration enhances episodic memory formation through neural oscillations. We presented participants with continuous auditory verbal and non-verbal narratives to memorise. Participants rated the perceived coherence of each narrative to provide a subjective measure of semantic elaboration. We assessed participants’ memory of the narratives depending on semantic elaboration and modality of encoding with a subsequent retrieval test. We recorded their electroencephalogram (EEG) to establish how semantic elaboration modulated the representation of narratives’ information proxied by neocortical alpha-beta oscillations during the encoding and retrieval. First, we found that semantic elaboration facilitated narrative information processing by speeding perceived coherence responses during encoding. Second, the magnitude of alpha-beta desynchronisation across the semantic network progressively increased with the chronological position of the events indexing information accumulation as the narratives unfolded. Third, the coherence scores of the narratives negatively correlated with the magnitude of alpha-beta oscillation desynchronisation later during successful retrieval. Our findings provide direct behavioural and alpha-beta oscillatory evidence of how semantic elaboration influences the time course of neural operations supporting information representation and reinstatement mediated by neocortical alpha-beta oscillations.

## INTRODUCTION

Episodic memory refers to rich memories associated with a unique spatiotemporal tag, while semantic memory represents one’s conceptual knowledge about words and objects (Renoult et al., 2019; Tulving, 2002). Although distinct, the interdependence of episodic and semantic systems to form memories has been long investigated since their seminal description by Tulving, but its neural basis still remains unclear (Greenberg & Verfaellie, 2010; Tulving, 1972). The *levels of processing* framework proposes that the degree (or depth) of semantic elaboration during episodic encoding determines the strength of a memory trace and its retention (Lockhart and Craik 1990; Craik & Tulving, 1975; Craik & Lockhart, 1972). This is because encoding an event is likely to trigger the activation of pre-existing semantic representations that link those elements. The more semantically congruent or coherent these elements are with each other, the more this activation facilitates the spread to broader, related semantic concepts during encoding, thereby promoting greater semantic elaboration, which in turn strengthens the memory trace (van Kesteren et al., 2012; Craik and Tulving, 1975; Schulman, 1974). Previous behavioural studies support this idea with participants better recalling words encoded within congruent events eliciting affirmative responses during categorisation than words embedded within incongruent events eliciting negative responses (Craik & Tulving, 1975 but see also Wagner et al., 1998; Lockhart and Craik 1990). Further studies also found that the semantic congruency between the different elements of an event typically improves subsequent memory, supposedly by promoting deep encoding (Wu & Hoffman, 2023; Quent et al., 2022; Tompary & Thompson-Schill, 2021; Amer et al., 2019; Frank et al., 2018; Packard et al., 2017; Bein et al., 2015; van Kesteren et al., 2013). In line with the *levels of processing* framework, functional magnetic resonance imaging (fMRI) evidence established that during encoding, activation in brain regions typically associated with semantic processing and semantic elaboration such as the Inferior Frontal Gyrus (IFG) and the lateral temporal lobes (Demirkan & Branzi, 2025; Branzi et al., 2020;2023; Jackson, 2021; Ralph et al., 2017; Badre et al. 2005; Wig et al., 2005; Wagner et al., 1998;), later predicts retrieval success of target words and associated features (Staresina et al., 2009). Further, the degree to which congruency between the elements of an episode enhances subsequent memory was positively correlated with increased encoding-related activation in the hippocampus, a key region involved in episodic memory encoding (van Kesteren et al., 2013; Staresina et al. 2009; for reviews Staresina & Davachi, 2008, 2006). Together, these results suggest that congruency prompts semantic elaboration, which in turn deepens encoding to form rich episodic memories.

Existing literature nevertheless presents limitations to fully address the influence of semantic elaboration on episodic encoding. Firstly, there is a lack of direct evidence for ‘enhanced semantic elaboration’ during encoding. The processing of congruent events should facilitate information processing as compared to incongruent events, leading to faster reading times or faster semantic judgement for instance. However, such effects have been rarely reported mostly because presenting words or sentences in isolation may not elicit different degrees of semantic elaboration during congruent *versus* incongruent event encoding in these studies (Staresina et al., 2009; Craik and Tulving 1975). In contrast, when semantic elaboration requires integrating new information with prior context during continuous encoding, participants process congruous narratives faster than incongruous ones in semantic tasks (Branzi & Lambon Ralph, 2023; Branzi et al., 2020), besides recalling more elements from short narratives when provided with a congruous compared to an incongruous context (Bransford & Johnson, 1972). These results suggest that the benefit of semantic elaboration is indeed dynamic and increases as semantic information accumulates over time during stimulus processing. Secondly, greater activations in brain regions typically associated to semantic processing are interpreted as evidence of enhanced semantic elaboration during the encoding of congruent *versus* incongruent events which in turn leads to better memory performance (Staresina et al., 2009). However, differences of activation in the inferior frontal gyrus (IFG) between trials referring to preexisting (congruous or plausible) versus non-preexisting (incongruous or not plausible) semantic associations may reflect differences in task difficulty or familiarity rather than differences of semantic coherence; see Demirkan & Branzi 2025; Jackson, 2021; Branzi et al., 2020). Thirdly, much of the evidence on semantic elaboration during encoding comes from slow fMRI, which does not allow to capture the fast time course of semantic processing and elaboration evidenced in other literature (Dirani & Pylkkänen, 2023; Rogers et al., 2021; Mollo et al., 2017; Clarke & Tyler, 2015; Miozzo, Pulvermüller, & Hauk, 2015). In contrast, electroencephalogram recordings (EEG) provide the necessary temporal resolution to investigate how neural oscillations orchestrate brain networks during speech-related encoding (Biau et al., 2025; Zioga et al., 2023; Lam et al., 2016). In particular, the desynchronisation of alpha-beta oscillations (8-30Hz) has been associated with access and representation of semantic information in the neocortex during memory encoding as well as retrieval (Branzi, Martin & Biau, 2023; Gastaldon et al., 2020; Roos and Piai, 2020; Hanslmayr, Staresina & Bowman, 2016; Piai et al., 2015; Hanslmayr et al., 2012; Jensen & Mazaheri, 2010). For instance, previous literature often reported an early desynchronisation followed by a progressive resynchronisation (or rebound) of neocortical alpha-beta oscillations during the successful retrieval of a memory (Griffiths et al., 2021; Pfurtscheller, Neuper, & Mohl, 1994). Further, elegant studies combining EEG to written verbal stimuli indeed conducted by Packard and colleagues evidenced modulations in the alpha-beta band during encoding associated with the positive effect of semantic congruence on subsequent memory (for instance Packard et al., 2020; 2017). However, how semantic elaboration influences the neural mechanisms mediating information representation through alpha-beta desynchronisation and boosts memory trace formation during sustained auditory encoding remains unclear.

The present study addresses the dynamic interplay between semantic elaboration and episodic encoding by overcoming these limitations. To mitigate the current pitfalls, we presented participants with continuous auditory verbal (spoken) and equivalent non-verbal (soundscape) narratives to memorise (**Figure 1A**). We prompted participants to process and integrate online information within an evolving context in different perceptual modalities to reveal if semantic processing in different perceptual modalities relies on similar or different oscillatory mechanisms. Importantly, participants rated the perceived semantic coherence of each narrative to provide a subjective measure of semantic elaboration. We assessed participants’ memory of the narratives depending on semantic elaboration and modality of encoding with a subsequent retrieval test (**Figure 1B**). We recorded the EEG of participants to establish how semantic elaboration modulated the representation of information of the narratives proxied by neocortical alpha-beta desynchronisation during the encoding and retrieval. We hypothesised (1) that semantic elaboration supports information processing during encoding by enhancing access to pre-existing semantic concepts. If so, narratives perceived as more coherent should be processed more quickly than less coherent ones, as reflected by faster response times for coherence judgments. This effect should be observed irrespectively of the perceptual modality. (2) If semantic elaboration promotes deeper encoding, then the degree of perceived coherence should predict subsequent memory performance. Therefore, details from coherent narratives should be better remembered than those from less coherent ones, regardless of perceptual modality. (3) If alpha-beta oscillations reflect dynamic semantic elaboration, then the magnitude of alpha-beta desynchronisation should progressively increase as the narratives unfold. This is because the incremental accumulation of semantic information should lead to increased concept activation and therefore prompt semantic elaboration, regardless of the perceptual modality of the narrative. If so, the degree of perceived coherence should predict alpha-beta desynchronisation magnitude across modalities as well. (4) Finally, if semantic elaboration promotes deep encoding via alpha-beta desynchronisation, then the perceived coherence should also influence information reinstatement later during memory retrieval and reflected by alpha-beta oscillation responses, independently from the narrative modality.

**Figure 1.**
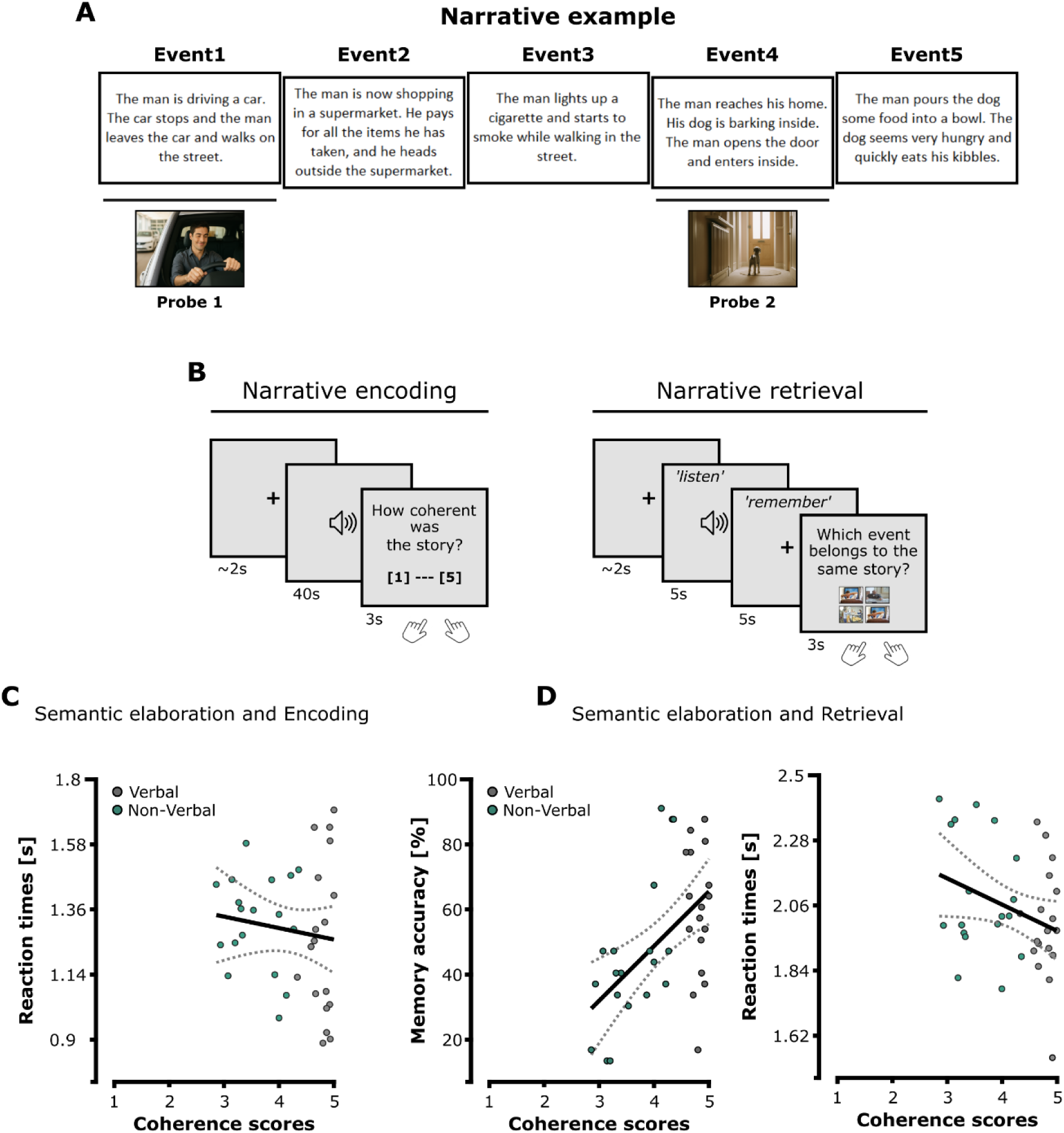
Memory task and behavioural performance. (**A**) Example of one narrative transcript composed of five sequential events presented either in verbal or non-verbal modality. The two probe 1 and probe 2 pictures were later used to test participants’ memory of the same narrative during the subsequent retrieval. Each picture depicted one key action or fact representing a specific event happening only in the cued narrative. For each trial of the retrieval, the correct picture was presented with three lures depicting events sourced from other narratives (note that here the two probe picture examples were computer-generated for anonymity purposes). (**B**) Overview of the memory task composed of the encoding and retrieval stages (the intermediate distractor stage is not included for simplification purposes). (**C**) Effect of semantic elaboration on coherence perception speed during the encoding of verbal and non-verbal narratives. (**D**) Effect of semantic elaboration on subsequent memory accuracy (left) and reaction times (right) during verbal and non-verbal narrative retrieval. For each scatter plot, the solid line represents the linear regression model and the dashed curves show 95% confidence intervals for the fitted regression line. The dots represent the data of each stimulus averaged across participants in the verbal (grey) and non-verbal (green) modalities (for complete visualisation of data, see **Supplementary Figure S1**).

## RESULTS

### Semantic elaboration modulates narrative encoding and enhances subsequent episodic memory (Hypotheses 1 & 2)

We designed a novel memory task consisting of an encoding, distractor and a retrieval stage (see **Figure 1A & 1B** and **Material**). During the encoding, participants attended and memorised verbal and non-verbal narratives composed of 5 sequential events, indicating their overall perceived coherence on a scale from 1 to 5 at the end of each presentation (See **VN1** and **NVN1** for equivalent verbal and non-verbal narrative examples). After a short distractor stage, participants’ memory of the narratives was assessed with a retrieval test. For each trial, participants were first auditorily cued with a short extract of one event previously encoded and were instructed to mentally retrieve its corresponding narrative. Participants then had to determine from four probe pictures displayed on the screen, which one depicted an event belonging to the narrative retrieved from the cue. The memory of each narrative was tested twice by using different cues and pictures. We tested our first and second hypotheses that semantic elaboration supports information processing during encoding by facilitating access to pre-existing semantic concepts and that this facilitation results in strengthened memory, independently from the perceptual modality. If so, the perceived coherence scores should predict participants’ speed to rate the narratives during encoding, as well as subsequent memory performance.

First, we fitted one separate linear mixed effects model using the reaction times of the ratings provided during the encoding task as the dependent variable to assess how perceived coherence impacted narrative processing (**Figure 1C**). The modality (verbal *vs.* non-verbal) and perceived coherence scores (1, 2, 3, 4 or 5) were used as two fixed effect factors at the level of the individual trials. By-subject information was also included as a random intercept in the model and the model was fitted using restricted maximum likelihood. Results revealed a significant effect of perceived coherence scores on reaction times (*F*(4, 481.86) = 6.486; *p* < 0.001), confirming that higher coherence scores inversely correlated with faster reaction times during narrative encoding (mean reaction times for the narratives with a perceived coherence score of 1: 1.941 ± 0.221 s; coherence score of 2: 1.353 ± 0.769 s; coherence score of 3: 1.376 ± 0.703 s; coherence score of 4: 1.35 ± 0.706 s; coherence score of 5: 1.147 ± 0.575 s). In contrast, the model revealed no effect of modality (*F*(1, 477.53) = 0.901; *p* = 0.333) or interaction between modality and perceived coherence scores on reaction times (*F*(3, 480.12) = 1.032; *p* = 0.378; mean reaction times for the verbal narratives: 1.223 ± 0.613 s and non-verbal narratives: 1.289 ± 0.686 s). Importantly, we controlled that the perceived coherence of verbal and non-verbal narratives reported by the participants during the encoding did not dependent on their modality (mean perceived coherence for the verbal narratives: 4.782 ± 0.583 and non-verbal narratives: 3.599 ± 0.107). We performed a linear regression model on the averaged coherence scores as well as reaction times across participants, between the narratives presented in verbal and equivalent non-verbal modalities (see **Supplementary Figure S2**). Results revealed a significant positive relationship for the perceived coherence scores across modalities, showing that verbal narratives with the highest scores were also highly rated in their non-verbal version (*F*(1, 16) = 4.922; *p* = 0.041; *R^2^* = 0.188). In contrast, no significant relationship was found for the reaction times (*F*(1, 16) = 0.559; *p* = 0.465; *R^2^*= - 0.027). This result confirms that perceived semantic coherence also captured the integration of information in the narratives, regardless of their modality of presentation. Narratives that were more difficult to semantic integrate and to represent in the verbal modality during encoding were also more difficult to represent in their equivalent non-verbal modality, potentially because of the specific spatiotemporal features and the sequence of their events.

Second, we fitted two separate linear mixed effects models using either the accuracy or reaction times of the retrieval task as the dependent variables to assess how perceived coherence predicted subsequent memory (**Figure 1D**). Modality (verbal *vs.* non-verbal) and coherence scores (1 to 5) from the encoding task were used as two fixed effect factors at the level of the individual trials, while by-subject information was included as a random intercept. Results revealed no significant effect of perceived coherence on subsequent memory accuracy (*F*(4, 761.81) = 1.597; *p* = 0.173). In contrast, it revealed a significant effect of modality (*F*(1, 816.84) = 18.849; *p* < 0.001), showing that participants better remembered verbal than non-verbal narratives (mean rate of correct responses for the verbal narratives: 0.602 ± 0.133 and non-verbal narratives: 0.426 ± 0.096). Crucially, the model also revealed a significant interaction between modality and perceived coherence (*F*(3, 797.88) = 3.947; *p* = 0.008). Bonferroni-corrected pairwise comparisons showed that memory accuracy of verbal narratives was more accurate compared to non-verbal narratives for narratives with a perceived coherence score of 2 (*p* = 0.005), 3 (*p* = 0.01) and 4 (*p* = 0.022) respectively. In the verbal modality, pairwise comparisons did not reveal differences of accuracy across the different coherence scores. In the non-verbal modality, pairwise comparisons showed that narratives with a coherence score of 2 were less retrieved than the ones with a score of 4 (*p* = 0.001) and 5 (*p* < 0.001). Importantly, we verified that participants retrieved memories of the narratives regardless of their modality of encoding by testing the overall mean accuracy against chance level (i.e., 0.25). To do so, we performed a one-sample T-test on the correct response rate for verbal and non-verbal trials together computed for each participant. Results confirmed that participants’ accuracy was significantly greater than chance level and that they successfully retrieved memories from the narratives (*t*(29) = 15.34388; *p* < 0.001; mean rate of correct responses for the verbal and non-verbal narratives: 0.514 ± 0.094). The second model did not reveal any significant effect of modality on reaction times (*F*(1, 500.55) = 1.396; *p* = 0.238), perceived coherence (*F*(4, 501.20) = 1.625; *p* = 0.167) or interaction between modality and perceived coherence (*F*(3, 503.62) = 0.371; *p* = 0.774). Mean reaction times for the verbal narratives: 2.023 ± 0.248 s and non-verbal narratives: 1.945 ± 0.242 s.

To summarise, participants successfully formed and retrieved meaningful memories from verbal and non-verbal narratives. Results confirm our first hypothesis that semantic elaboration facilitates information processing during encoding, which was reflected by speeding the perceived coherence judgment independently from the narrative modality. In contrast, these results partially support our second hypothesis that semantic elaboration during encoding enhances subsequent memory similarly across modalities because this effect was observed in the non-verbal modality only. Furthermore, verbal narratives were better remembered than non-verbal ones, regardless of the perceived coherence scores.

### Alpha-beta desynchronisation reflects the progressive accumulation of information to support semantic elaboration during encoding (Hypothesis 3)

We then addressed our third hypothesis that dynamic semantic elaboration increases with the accumulation of integrated information as narratives unfolds. Accordingly, we expected a progressive alpha-beta oscillation desynchronisation in brain regions associated with semantic processing (**Figure 2A & 2B**).

**Figure 2.**
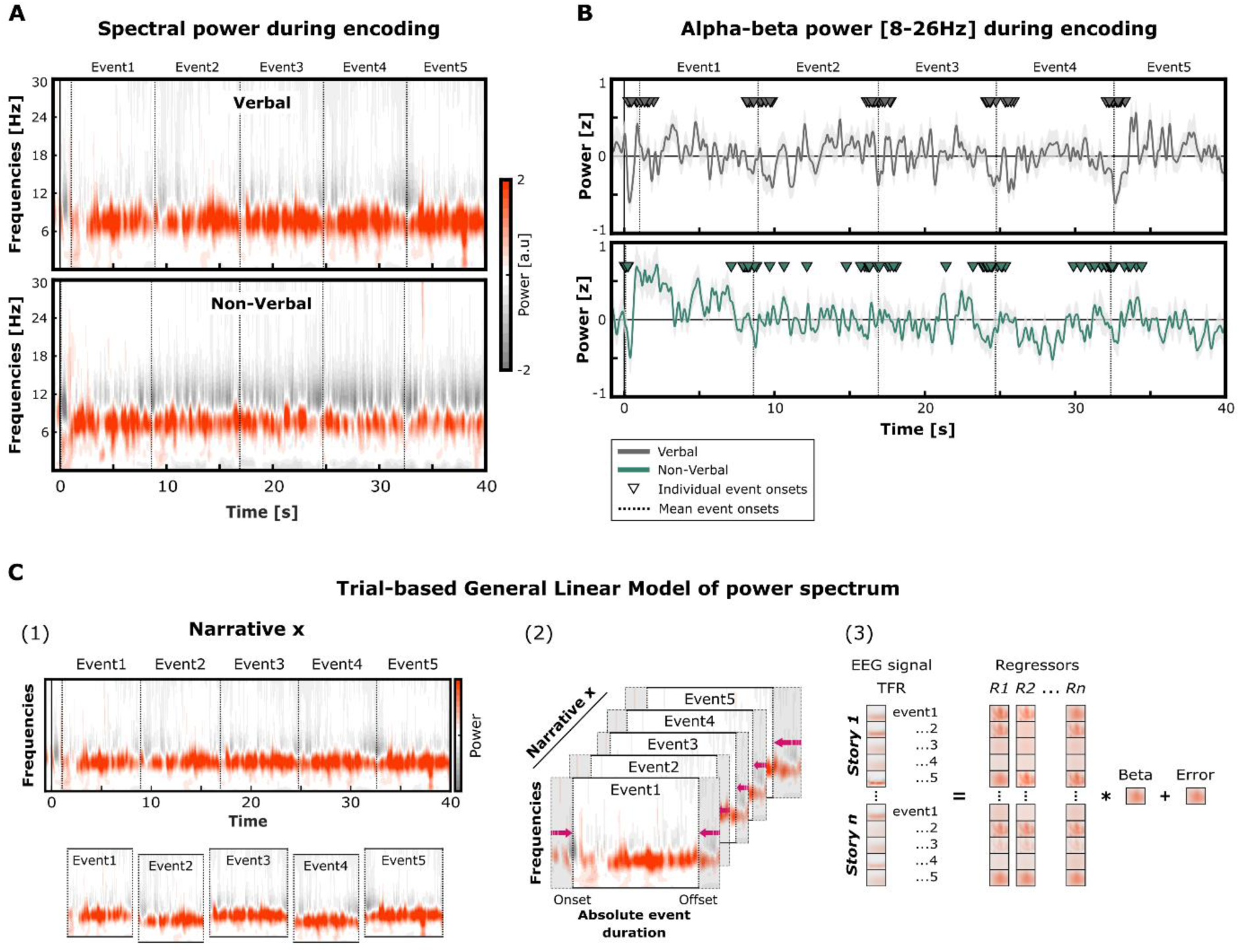
Power spectrum changes during narrative encoding. (**A**) Time-frequency representation of power spectrum when participants encoded narratives. (**B**) Time course of averaged alpha-beta power [8-26Hz] during narrative encoding (solid line: mean power; grey area: standard deviation). The magnitude of alpha-beta power desynchronisation around event onsets increased as measure as the narrative unfolded, i.e., with the chronological position of the event indexing information accumulation. (**C**) Trial-based general linear model was used to quantify the effect of modality, event position and their interaction on power spectrum during narrative encoding.

We first estimated how the chronological position of the sequential events (from the 1^st^ to 5^th^ event) reflects accumulation of semantic content and predicts changes in the power spectrum in both modalities, by using a trial-based regression model (see **Figure 2C** and **Material**). (1) each continuous power spectrum evoked during verbal and non-verbal encoding was segmented within the sequential events from 1 to 5, an ordered in a single-trial spectral power fashion. (2) As the event durations varied within and across narratives, each event spectrum was interpolated in the time dimension to fit a generic time-frequency matrix (1-30 Hz; 126 time-points). (3) The power spectrum was estimated using a general linear model including five regressors: (i) the modality of the narratives, (ii) the chronological position of each event within the narratives, (iii) the interaction between modality and position of events, (iv) the perceived coherence score of the narrative and (v) the Word2vec coherence scores as regressors. While the first three regressors (i, ii and iii) served to address our hypothesis, the perceived coherence and Word2vec coherence regressors (iv and v) reflected semantic parameters to account for participant- and event-related variance. A beta coefficient for each regressor was obtained for every frequency, time-point, and channel, and pooled across participants for statistical assessment by means of cluster-based permutation tests. Cluster-based permutations performed on the regressor coefficients revealed a significant negative cluster for the interaction between modality and event position during narrative encoding (*p* = 0.0045, cluster size = -1.531x10^4^, mean t-statistic within cluster = -2.596; Cohen’s *d_z_* = -0.474). This effect was found through the event duration within narrative and broadly across central regions of the scalp (**Figure 3A)**. In contrast, no significant cluster was found for the modality or event position regressors on the power spectrum (no significant cluster was found for the perceived coherence score regressor, but non-essential results were found for the Word2vec coherence score, see **Supplementary Figure S3**).

**Figure 3.**
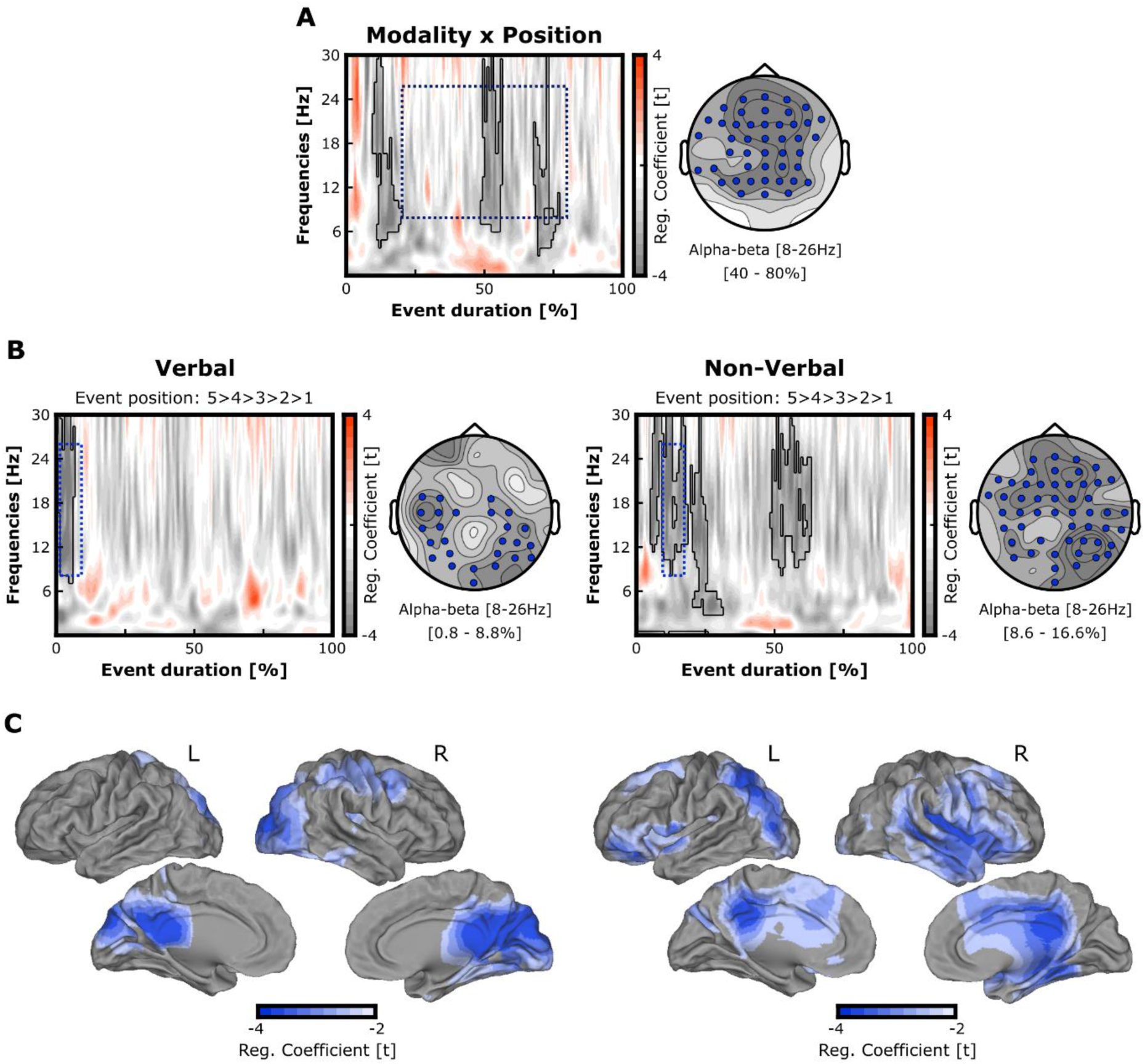
Alpha-beta activity progressive desynchronisation with the accumulation of semantic information during narrative encoding. (**A**) Time-frequency representation and topography of the power spectrum changes induced by the interaction between the modality and events’ position during narrative encoding. The time-frequency plots the t-values averaged across the significant channels (indicated by blue dots on the topography), and the black line outlines significant t-values. The topography shows the t-values for the 8-26Hz alpha-beta activity within the central portion of the event duration, i.e., [20 - 80%] and indicated by the blue dashed rectangle. (**B**) Time-frequency representations and topographies of the power spectrum changes induced by the position of the events during verbal and non-verbal narrative encoding. Alpha-beta oscillations desynchronized with online semantic processing during encoding (evidenced by the gradual darkness of the grey colour in the plots). The time-frequency representations plot the t-values averaged across the significant channels (indicated by blue dots on the topographies), and the black line outlines significant t-values. The topographies show the t-values for the 8-26Hz alpha-beta activity within the event portions of interest (blue dashed rectangles) for verbal and non-verbal narratives, respectively [0.8 - 8.8%] and [8.6 - 16.6%]. (**C**) Source localisation of the alpha-beta event position effect in verbal and non-verbal narratives, within the time-windows of interest evidenced by the blue dashed rectangle in A (threshold at significant t-values).

The negative cluster observed for the interaction effect may potentially reflect redundancy and common information representation across brain activities supporting semantic processing (Gelens et al., 2024). We then addressed whether similar mechanisms took place during the encoding of both verbal and non-verbal narratives. We estimated the effect of events’ position on the power spectrum during encoding using two separate trial-based regression models for verbal or non-verbal narratives (**Figure 3B and 3C**). To do so, the separate power spectrums in verbal and non-verbal modalities were estimated using a general linear model including the position of each event as the regressor of interest, and the perceived coherence score of the narratives as well as the Word2vec coherence scores of events as semantic regressors to account for participant and stimulus variance. A beta coefficient for the regressors was obtained for every frequency, time-point and channel, and pooled across participants for statistical assessment by means of cluster-based permutation tests. Statistical assessment was only performed for the regressor of interest (i.e., event position). The *p* values of the significant clusters were Bonferroni-corrected for multiple comparisons across the modalities. Cluster-based permutations performed on regressor coefficients at scalp level (**Figure 3B**) revealed a significant negative cluster predominantly localised in the alpha-beta range [8-26Hz] of the power spectrum during the encoding of both verbal (*p* = 0.028, cluster size = -1.837x10^4^, mean t-statistic within cluster = -2.578; Cohen’s *d_z_* = -0.471) and non-verbal narratives (*p* = 0.010, cluster size = -2.836x10^4^, mean t-statistic within cluster = -2.608; Cohen’s *d_z_* = -0.476). In both cases, alpha-beta oscillations desynchronised more as the event occurred later in the narrative, with this modulation taking place early after the events’ onset. Significant alpha-beta desynchronisation was observed in the bilateral centro-occipital regions across both narrative modalities. The source analysis of the alpha-beta event position effect (**Figure 3C**) revealed a significant negative cluster in the bilateral occipital regions for verbal (*p* = 0.012, cluster size = -1.931x10^3^, mean t-statistic within cluster = -2.791; Cohen’s *d_z_* = -0.51) and non-verbal narrative encoding (*p* < 0.001, cluster size = - 3.384x10^3^, mean t-statistic within cluster = -2.92; Cohen’s *d_z_* = -0.533), confirming the correlation between the event chronological position and the magnitude of alpha-beta desynchronisation at source level. The effect of event position on alpha-beta desynchronisation extended to frontal, parietal and temporal lateral regions that are typically associated to semantic processing (Demirkan & Branzi, 2025; Jackson, 2021). Altogether, these results support our hypothesis that the accumulation of integrated information prompts semantic elaboration as narratives unfolds, which induces a progressive alpha-beta oscillation desynchronisation providing access to pre-existing concepts. Further, results suggest that similar neural mechanisms account for semantic elaboration regardless of the modality of the narratives (Packard & Soto-Faraco, 2025; Tulving & Madigan, 1970).

### Semantic elaboration shapes information reinstatement during successful retrieval (Hypothesis 4)

According to our fourth hypothesis, semantic elaboration impacts neural operations mediating information representation during narrative encoding, and therefore should determine its reinstatement during subsequent memory retrieval. If so, perceived coherence scores during encoding should influence alpha-beta desynchronisation during successful retrieval and regardless of the perceptual modality. To address our hypothesis, we then estimated the power spectrum during retrieval with a trial-based regression model by including the regressors modality (verbal or non-verbal) and perceived coherence score of the narrative during encoding (from 1 to 5). Only successful trials (i.e., hits) were included. Permutation analysis performed on the regressor coefficients revealed a significant positive effect of perceived coherence on power spectrum in the alpha-beta range, extending to the theta range, and localised in the fronto-centro-parietal regions (*p* = 0.004, cluster size = 1.157x10^4^, mean t-statistic within cluster = 2.543; Cohen’s *d_z_* = 0.464; Figure 4). Source analysis in the significant time-window [2.8 - 3.4s] confirmed a positive correlation between perceived coherence and alpha-beta power in bilateral centro-parietal areas (*p* = 0.019, cluster size = 491.069, mean t-statistic within cluster = 2.468; Cohen’s *d_z_* = 0.45; **Figure 4)**. The model also revealed an effect of modality on the power spectrum with a significant cluster (*p* = 0.005, cluster size = 1.294x10^4^, mean t-statistic within cluster = 2.546; Cohen’s *d_z_* = 0.465; see **Supplementary S4**). As the behavioural results revealed a significant interaction between modality and perceived coherence, we initially aimed to include the interaction regressor in the model to assess its effect on the power spectrum during successful retrieval as well. However, due to the limited variance in perceived coherence ratings provided by participants for the verbal narratives, the model suffered from multicollinearity and rank deficiency, which prevented us from including this regressor, as we did during encoding.

**Figure 4.**
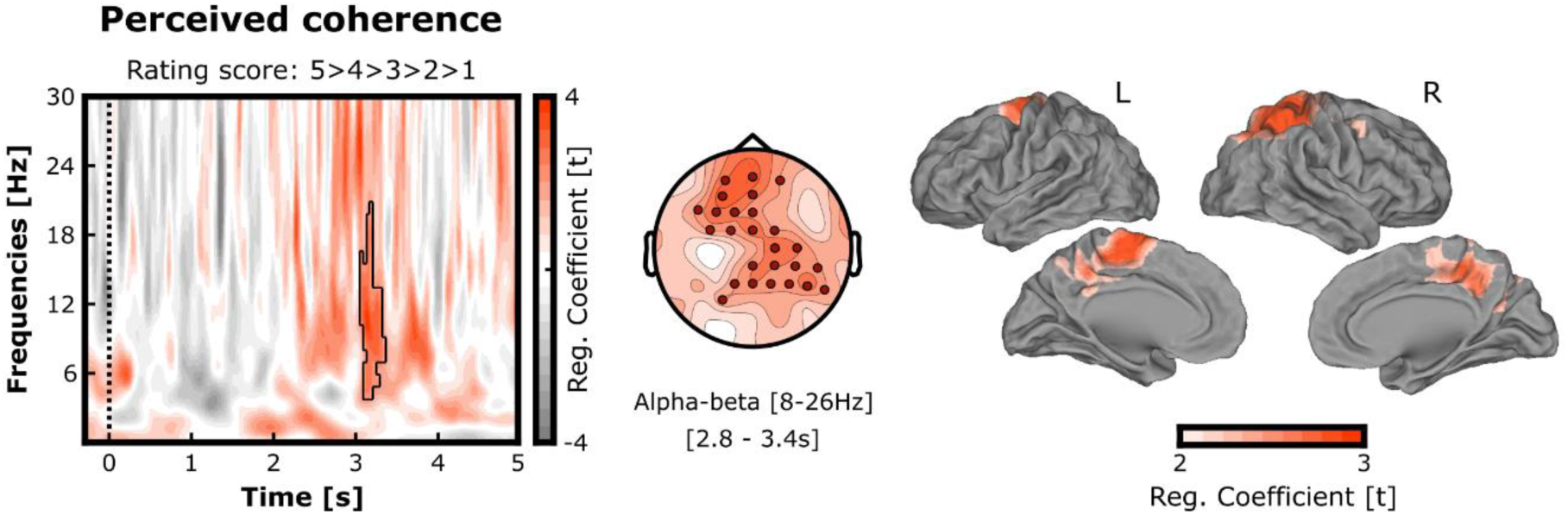
Effect of perceived coherence on alpha-beta oscillations during successful retrieval. Left and centre: Time-frequency representation and topography of the effect of perceived coherence on power spectrum during memory reinstatement (verbal and non-verbal narratives collapsed together). Alpha-beta desynchronisation decreased with semantic elaboration level estimated by the perceived coherence scores provided during encoding (evidenced by the gradual darkness of the red colour in the plots). The time-frequency representation plots the t-values averaged across the significant channels (indicated by red dots on the topographies), and the black line outlines significant t-values. The topography shows the t-values for alpha-beta activity (8-26 Hz) within the time-window of interest [2.8 - 3.4s] outlined by the black line in the time-frequency plot. Right: Source localisation of the effect of perceived coherence on alpha-beta oscillations in the time-window of interest (threshold at significant t-values).

For completeness, we also conducted analysis on the time course of alpha-beta power modulation during successful memory retrieval of verbal and non-verbal narratives. Results confirmed a similar early desynchronisation followed by a progressive resynchronisation (or rebound) of neocortical alpha-beta oscillations, which was expected from previously literature (Griffiths et al., 2021; Pfurtscheller, Neuper, & Mohl, 1994). The power decrease occurring around 500 ms after the onset has indeed been described and hypothesised to proxy the reinstatement of neurophysiological computation to represent the information retrieved from a memory (see **Supplementary Figure S5**).

Our results confirmed our fourth hypothesis that semantic elaboration not only influences the time course of information representation during narrative processing but also its reinstatement during successful retrieval, mediated by the desynchronisation of neocortical alpha-beta oscillations. Second, the modality of the narrative during encoding determined differences in alpha-beta desynchronisation amplitude that were reflected by memory performance during subsequent retrieval, i.e., verbal narratives were better remembered and their recollection was accompanied by an attenuation of alpha-beta desynchronisation compared to non-verbal narratives.

## DISCUSSION

The present study investigates for the first time the neural mechanisms accounting for the dynamic interplay between semantic and episodic memory during the encoding of naturalistic narratives. Our work goes beyond previous studies in different ways: First, we provide direct behavioural and oscillatory evidence of how semantic elaboration influences meaningful information representation during encoding and retrieval stages of narrative memories. Second, our findings evidence the cross-modal benefit of semantic elaboration to encode verbal and non-verbal narratives, regardless of the perceptual modality.

The results confirmed our first hypothesis that semantic elaboration speeds up the processing of information because participants rated narratives more rapidly as their perceived coherence increased during encoding. This aligns with the *levels of processing* framework predicting that semantic congruence within an event broadens the spread of pre-existing representation activations, which facilitates the integration of information (Lockhart and Craik 1990; Craik & Tulving, 1975; Craik & Lockhart, 1972). Here, participants were prompted to integrate new information with a prior context as a narrative unfolded to generate and continuously update a meaningful representation throughout stimulus presentation. Accordingly, the integration of incoming information closely relating to the prior parts of the story triggered a wider range of overlapping semantic representations -or *schemas*- of their relationship, compared to less congruent elements (Tibon, Cooper & Greve, 2017; Packard et al., 2017; van Kesteren et al., 2012). A broader spread of pre-existing schema activations deepened the level of details in the narrative representation and speeded the final coherence scoring decision (Branzi & Lambon Ralph, 2023; Branzi et al., 2020). Importantly, the coherence scores predicted rating speed for both verbal and non-verbal narratives, supporting our hypothesis that semantic elaboration enhances the activation of pre-existing high-level concepts rather than modality-specific features (Tulving & Madigan, 1970). For instance, Packard and Soto-Faraco (2025) recently showed that incidental picture encoding accompanied by congruent sounds led to better long-term memory than incongruent pairs, illustrating the cross-modal benefit of semantic elaboration.

Our second hypothesis that semantic elaboration enhances episodic memory independently from the modality was partially supported by the retrieval performance. As expected, participants successfully retrieved memories from both verbal and non-verbal narratives, but coherence during encoding predicted subsequent memory accuracy in the latter case only. Previous studies have already established that semantic congruence promotes encoding of written, visual, or multisensory events, such that congruent items are later remembered better than incongruent ones (Packard & Soto-Faraco, 2025; Packard et al., 2020; 2017; van Kesteren et al., 2013; Atienza, Crespo-Garcia & Cantero, 2011; Staresina et al. 2009; Craik and Tulving, 1975; Schulman, 1974). Here, we demonstrate that congruence during encoding facilitates episodic memory for long auditory stories as well, in particular for soundscape narratives in which listeners relied on semantic knowledge to interpret sounds and generate meaningful representations of sequential events. Notably, prior research has almost exclusively focused on isolated events, such as direct congruence between pairs of stimuli or a categorial cue. To our knowledge, the present study is the first to investigate semantic elaboration during sustained narrative processing, in which congruence was established through the integration of accumulating information with an evolving context and pre-existing individual knowledge (e.g., *“Are these new elements normally expected to occur within a sequence of congruent events, given the previous part of the narrative and my prior experience?”*). Importantly here, participants determined congruency of the narratives themselves in order to promote explicit semantic processing during encoding and to target memory before consolidation or retrieval (Packard et al., 2017). Further, it allowed us to consider interindividual differences of personal knowledge contributing to semantic elaboration among participants (Tibon, Cooper & Greve, 2017; Packard et al., 2017). The lack of behavioural evidence for verbal narratives may be explained by two non-exclusive possibilities. First, verbal narratives were almost exclusively scored as highly coherent, which may have prevented the detection of a congruence-memory relationship due to insufficient variability. Alternatively, verbal memories may have been too easily retrieved to capture the cross-modal benefit of semantic elaboration, as auditory words are better remembered than equivalent sounds in serial recall tasks (Paivio, Philipchalk & Rowe, 1975; Philipchalk & Rowe, 1971). It might be that semantic elaboration yields incremental benefits for encoding only up to a certain point, beyond which it ceases to be critical for successful memory. The fact that memory accuracy between verbal and non-verbal modalities did not differ anymore for narratives with the highest coherence scores of five supports this interpretation. Future research should investigate this relationship using semantic coherence measures sampled at multiple time points during encoding to determine whether continuous measures during encoding better predict changes in oscillatory patterns.

Our EEG results support the third and fourth hypotheses that semantic elaboration shapes the neural mechanisms mediating information representation and reinstatement reflected by the desynchronisation of alpha-beta oscillations (Roos and Piai 2020; Hanslmayr et al., 2012; Hanslmayr, Staresina & Bowman, 2016; Piai et al., 2015; Jensen & Mazaheri, 2010). We first found that the magnitude of alpha-beta desynchronisation progressively increased with the chronological position of the events indexing information accumulation as the narratives unfolded. Previous literature suggested that the desynchronisation of alpha-beta oscillations reflects access to semantic information across the neocortex networks. According to the *levels of processing* framework, encoding the narratives prompted semantic spread to continuously integrate incoming information with previous context and to update an online representation of semantic content. Here we assume that the dynamic representation of semantic information was enabled by the gradual loosening of alpha-beta synchronisation, which typically functions as a “gatekeeper,” preventing knowledge from becoming labile (Jensen & Mazaheri, 2010; Lockhart and Craik 1990; Craik & Tulving, 1975; Craik & Lockhart, 1972). Importantly, alpha-beta desynchronisation and event positions correlated in brain regions associated with semantic processing, including the IFG as well as bilateral occipito-parietal and temporal regions (Branzi et al., 2023; Branzi et al., 2020; Gastaldon et al., 2020; Paunov, Black & Fedorenko, 2019;). To our knowledge, we provide the first neurophysiological evidence of semantic elaboration influence over the time course of information processing, and mediated by alpha-beta oscillations across the “semantic network” (Jackson 2021; Ralph et al., 2017).

During successful retrieval, semantic elaboration influenced neural mechanisms mediating information reinstatement as well, and we found that coherence scores of the narratives negatively correlated with the magnitude of alpha-beta oscillation desynchronisation. It is possible that greater semantic spread during narrative processing enhanced memory encoding and later facilitated it recollection, resulting in reduced alpha-beta desynchronization during retrieval. A previous study also reported a pattern of increasing encoding-related activity in the memory regions of the MTL with decreasing congruency in object-scene associations (van Kesteren et al., 2013). Here, this association was observed in the mid cingulate, as well as parietal regions such as the precuneus and cuneus, which have been associated both with episodic and semantic memory retrieval (Tanguay et al., 2023; Baldassano et al., 2017). Their engagement could reflect the recruitment of neural mechanisms supporting the reinstatement of information previously elaborated during the encoding, and recruited while participants recollected memories (Palacio & Cardenas, 2019). Encoded-related activity in the cingulate cortex has indeed been shown to increase with the congruency between the pairs of stimuli (van Kesteren et al., 2013), which aligns with our results reporting a link between semantic congruency and subsequent retrieval. Other works showed that the magnitude of alpha-beta desynchronisation increased with the complexity of the retrieval task such as correct rejection, recognition or lure discrimination (Karlsson et al., 2023; Karlsson et al., 2020). Our inverse correlation between perceived coherence and alpha-beta activity captured the degree of difficulty to reinstate narrative information that was already difficult to represent during previous encoding, due to stimulus-specific features.

Interestingly, the relationship between event position, perceived coherence and alpha-beta desynchronisation found during the encoding-retrieval stages was also observed in the posterior cingulate cortex (PCC) and the precuneus for both verbal and non-verbal narratives. These regions have been shown to respond to the coherence of events’ structure (Simony et al., 2016; Lerner et al., 2011; Hasson et al., 2008), and recent studies have associated the PCC with the construction of complex semantic “gist-like” representations across auditory and visual modalities (Branzi & Lambon Ralph 2022; Baldassano et al., 2017). Reflecting our behavioural results, the overlap of alpha-beta desynchronisation across verbal and non-verbal encoding indeed suggests that semantic elaboration facilitates encoding and later influences retrieval regardless of the perceptual modality (Tulving & Madigan, 1970).

## CONCLUSION

The present study addresses how semantic processing dynamically interacts with episodic memory to facilitate the encoding of verbal and non-verbal narratives. For the first time, our results demonstrate that the semantic elaboration influences the time course of the neural operations supporting information representation and reinstatement in the brain, mediated by alpha-beta oscillation desynchronisation across the neocortex. Further, our findings suggest that semantic elaboration facilitates memory formation and retrieval of both verbal and non-verbal narratives, regardless of the perceptual modality.

## SUPPLEMENTARY INFORMATION

Supplementary information contains five figures.

## AUTHOR CONTRIBUTION

E.B and F.B designed the study and collected the data. E.B and F.B analysed the data. E.B and

F.B wrote the paper. The authors discussed the results and commented on the manuscript.

## ACKNOWLEDGEMENT

This work was supported by a Sir Henry Wellcome Fellowship (210924/Z/18/Z) and a Tenure-Track Fellowship funded by the University of Liverpool awarded to E.B.

## KEY RESOURCES TABLE

| REAGENT or RESOURCES | SOURCES | IDENTIFIER |
| --- | --- | --- |
| **Software and algorithms** |  |  |
| MATLAB | www.mathworks.com | R2023 |
| R | https://www.r-project.org/ | 4.5.1 |
| lme4 package | https://cran.r-project.org/web/packages/lme4/index.html | 1.1-37 |
| lmerTest package | https://cran.r-project.org/web/packages/lmerTest/index.html | 3.1-3 |
| Psychophysics Toolbox | psychtoolbox.org | 3 |
| Fieldtrip | www.fieldtriptoolbox.org | 20230427 |
| SPM | Wellcome Trust Centre for Neuroimaging | 12 |
| Caret | Washington University School of Medicine | 5.65 |
| ER-3C system | www.etymotic.com | EEG compatible earphones |

## RESOURCE AVAILBILITY

Lead contact

Further information and requests for resources and reagents should be directed to and will be fulfilled by the lead contacts Emmanuel Biau or Francesca Branzi.

Material availability

This study did not generate new unique reagents. Data and code availability

The data sets and original codes have been deposited at https://osf.io/xxxxx/ and will be made available after publication. Additional requests for information related to the present article will be directed to Emmanuel Biau and Francesca Branzi.

## EXPERIMENTAL MODEL AND SUBJECT DETAILS

Thirty-two healthy participants took part in the experiment. All participants were English native speakers and right-handed. All of them reported normal or corrected-to-normal vision and hearing. All participants received financial reimbursement for taking part in the experiment. Two participants were excluded because of excessive artifacts in the EEG data. This left thirty participants (mean age = 23.47 years ± 4.14; 20 females) for data analysis. All participants gave written informed consent. Ethical approval was granted by the University of Liverpool Research Ethics Committee complying with the Declaration of Helsinki.

## METHOD DETAILS

### Auditory narrative stimuli

Eighteen pairs of narratives presented in verbal or non-verbal modalities were created for the present study (**Figure 1A**). The verbal stimuli consisted of 40 seconds(s)-long spoken stories narrated by a native English male speaker and composed of five chronological ‘events’ lasting approximately 8 s each (mean duration 1^st^ event = 7.87 ± 0.76 s; 2^nd^ event = 8.00 ± 0.83 s; 3^rd^ event = 7.85 ± 0.90 s; 4^th^ event = 7.82 ± 0.66 s; 5^th^ event = 7.42 ± 0.39 s). Each event corresponded to a part of the story where new information about characters, locations or occurrences with respect to the preceding event was introduced. The corresponding non-verbal stimuli consisted of soundscape stories of the same length, describing the same events as in the verbal version also lasting approximately 8 s (mean duration 1^st^ event = 8.51 ± 1.17 s; 2^nd^ event = 8.3 ± 1.73 s; 3^rd^ event = 7.80 ± 1.04 s; 4^th^ event = 7.66 ± 1.54 s; 5^th^ event = 7.63 ± 1.17 s). Non-verbal stimuli were created using environmental natural sounds sourced from different websites (e.g., https://freesound.org/ or https://sound-effects.bbcrewind.co.uk/) and edited using Adobe Audition. All verbal and non-verbal narratives were edited and exported in *.mp3* format (44100 Hz sampling rate, mono) using Shotcut (Meltytech, LLC).

### Between-event semantic relatedness within narratives

For each narrative, we measured the between-event semantic relatedness using Word2vec modelling (Mikolov et al., 2013) to reflect event-related variance across stimuli when estimating the power spectrum in our general linear models. Semantic relatedness measures were obtained by subjecting narrative events to computational analyses based on a pre-trained Google model (https://code.google.com/archive/p/word2vec/). In essence, Word2vec provides vector-based representations of words’ meaning combined linearly to represent the meaning of each event composing the narratives (Hoffman, 2019): (1) For every verbal narrative, we obtained five vectors representing the semantic content of each event using their individual text transcripts with the pre-trained Google model. (2) Within each narrative, the vector of an event *n* was compared to the vector combining the prior events from 1 to *n*- 1, by using a cosine similarity metric. The cosine estimates their similarity with a unique value comprised between zero and one, i.e., one indicating that the semantic representation of an event is closely related to the prior context and zero indicating a drop of relative semantic relatedness. (3) The same operation was repeated by comparing the vector of the next event *n+1* with the vector combining the prior events from 1 to *n*, and repeated until the last event of the narrative. We therefore obtained five semantic relatedness values per narrative in the chronological order of the events (for further details, see Branzi & Lambon-Ralph, 2022). The semantic relatedness values were determined with verbal narratives and served as a semantic parameter controlling for event-related variance for equivalent non-verbal narratives as well. We did so because the specific spatiotemporal features and event sequences were aligned within each pair of narratives in the verbal and non-verbal versions, which was confirmed with a positive correlation of perceived coherence scores between verbal and non-verbal narratives during encoding (i.e., verbal narratives that were harder to integrate during encoding were also harder to integrate in the corresponding non-verbal modality)

### Experimental procedure and tasks

Participants seated comfortably in the testing room approximatively 60 cm away from the screen. The task was programmed with Matlab (R2021a; The MathWorks) and Psychophysics Toolbox-3 (Brainard & Vision, 1997). The sound stimuli were presented between 70 and 80 dB through EEG-compatible insert earphones (Etymotic Research, Elk Grove Village, IL) while participants visualised a fixation cross on a middle-grey screen (1920x1200 pixel resolution). The procedure consisted of three stages: (1) encoding, (2) distractor and (3) retrieval (**Figure 1B**). The whole session, including the set-up of the EEG cap, lasted for 120 minutes. Regular self-paced breaks were allowed for participants to rest.

### Encoding Task

During the encoding stage, each participant was presented with eighteen narratives—nine verbal and nine non-verbal—each presented only once. To avoid priming effects, for any given participant, verbal and non-verbal stimuli did not share the same semantic content. However, across all participants, both the verbal and non-verbal versions of each narrative were presented. The order of the narratives was randomised for each participant, and the version (verbal or non-verbal) of each narrative presented during the encoding was counterbalanced across participants. Each trial started with a red fixation cross (jittered duration: 2000 - 2500 ms) to indicate the start of the audio stimulus followed by the auditory presentation of a narrative. During the narrative presentation, a grey fixation cross was displayed on the screen to minimise eye movement contamination of the EEG signal. Participants were instructed to carefully attend to the auditory stimuli and try to understand their content as the narratives unfolded. Participants were also instructed to memorize the stimuli for the subsequent memory test taking place later. At the end of each narrative, participants rated its coherence with the keyboard as follows: “1” = don’t know, “2” = very incoherent, “3” = somewhat incoherent, “4” = somewhat coherent, “5” = very coherent. Participants were given a time limit of three seconds to indicate their perceived coherence score. The next trial began immediately after the participant’s response or when the time limit elapsed.

### Distractor Task

Following the encoding stage, participants completed a short distractor task of twenty trials. The distractor task aimed to prevent lingering working memory effects from the encoding to the retrieval stages. Each trial started with a brief fixation-cross (jittered duration: 1000 - 1500 ms) followed by the presentation of a random number (from 1 to 99) displayed at the centre of the screen. Participants had to determine as fast and accurately as possible whether this number was odd or even by pressing “R” (“odd”) or “P” (“even”) on the keyboard.

### Retrieval Task

After the distractor task, the participants performed the retrieval task to assess their memories of the narratives. Each trial started with a brief fixation-cross (jittered duration: 2000 - 2500 ms) followed by the presentation of an auditory cue. The auditory cue consisted of a meaningful 5s segment extracted from one of the events in a previously encoded narrative. Participants were instructed to listen (“Listen”) to the auditory cue and were given additional 5s to silently recall the corresponding narrative (“Remember”). Then, four pictures appeared on the screen and participants had to indicate as fast and accurately as possible which one depicted an event belonging to the narrative retrieved with the auditory cue (four-alternative-forced-choice task; one correct option and three distractors). Memory for each narrative was tested twice but using different cues and picture options. The retrieval task included a total of 36 trials per participant. Participants had a time limit of 3 s to respond using the keyboard, by pressing keys “1” to “4” to indicate pictures 1 to 4). The next trial began immediately after the participant’s response or when the time limit elapsed. The position of the event represented by the probe pictures varied across narratives to control for event position effect on memory recollection.

### EEG data analysis

### EEG acquisition and preprocessing

Continuous EEG signal was recorded using a 64 channel BioSemi ActiveTwo system (BioSemi, Amsterdam, Netherlands). Vertical and horizontal eye movements were recorded from additional electrodes placed approximatively one cm to the left of the left eye, one cm to the right of the right eye, and one cm below the left eye. Online EEG signals were digitalized using BioSemi ActiView software at a sampling rate of 512 Hz. EEG data were preprocessed offline using Fieldtrip (Oostenveld et al., 2011) and SPM 12 (Wellcome Trust Centre for Neuroimaging). Continuous EEG signals were bandpass filtered between 1 and 120 Hz and bandstop filtered (48-52 Hz and 98-102 Hz) to remove line noise at 50 and 100 Hz. Data from encoding and retrieval stages were epoched from 2000 ms before stimulus onset to 2000 ms after stimulus offset. Data from the distractor stage were not analysed. Trials and channels with artefacts were excluded by visual inspection before applying an independent component analysis (ICA) to remove components related to ocular artefacts. Excluded channels were then interpolated using the method of triangulation of nearest. After re-referencing the data to an average reference, the remaining trials with artefacts were manually rejected through a final visual inspection (on average, 5.13 ± 5.35 trials were rejected per participant).

### Time-frequency decomposition at sensor level

Time-frequency decomposition was computed on the encoding and retrieval epochs of verbal and non-verbal modalities at the sensor level. For the encoding trials, data were first convolved with a 5-cycle Morlet wavelet (from 1 to 40 Hz; 1 Hz step and 40 ms time steps) from -2 s to 40s with respect to narratives’ onset. For the retrieval trials, data were processed similarly from -2 s to 5 s with respect to “remember” period when participants successfully recollected memories from the auditory cue (hits only). Second, background fractal activity was attenuated in the time-frequency representation (TFR) by subtracting 1/f characteristic from the power spectrum using an iterative linear fitting procedure (Griffiths et al. 2021; Biau et al., 2022). This step generated two vectors: one vector containing the values of each wavelet frequency A, and another vector containing the power spectrum for each electrode-sample pair B. Both vectors were then put into log-space to approximate a linear function to get the slope and intercept of the 1/f curve. The linear equation Ax = B was resolved using least-squares regression, where x is an unknown constant describing the curvature of the 1/f characteristic. The 1/f fit Ax was then subtracted from the log-transformed power spectrum B.

### Time-frequency decomposition at the source level

The preprocessed data were reconstructed at source level by using a standard head model and MRI scan templates from Fieldtrip. The electrode positions on the scalp were defined using a template from Fieldtrip to prepare the source model. Time-locked EEG data were reconstructed applying a linearly constrained minimum variance (LCMV) beamforming approach implemented in Fieldtrip (van Veen et al., 1997). For each participant, time-series data were reconstructed for 2020 virtual electrodes. Time-frequency decomposition and trial-based multiple regression analyses were conducted at source level as for scalp level. The maximum voxel activation regions were defined by using the automated anatomic labelling atlas (AAL; Tzourio-Mazoyer et al., 2002).

### Statistical Analysis Behavioural statistical analysis

### Encoding

To assess whether semantic elaboration supports information processing during encoding by facilitating access to pre-existing semantic concepts, and strengthens memory, we computed individual coherence scores provided by participants for each stimulus during encoding, along with their corresponding reaction times. Narratives for which participants did not respond within the time limit were labelled as “misses”. We then fitted a separate linear mixed effects model using the reaction times in the encoding task as the dependent variable, and modality (verbal and non-verbal) as well as the coherence scores (1, 2, 3, 4 or 5) as two fixed effect factors at the level of the individual trials (LME; Baayen, Davidson, & Bates, 2008). By-subject random intercept was included in the model to account for variability in the reaction times at the level of individual participants. The model parameters were estimated using the restricted maximum likelihood approach. The modelling was performed using R (R Core Team, 2025 version 4.5.1) and the “lme4” package within R (Bates et al., 2015). To obtain the significance *p*-values using Satterthwaite’s method, we used the “lmerTest” package (Kuznetsova et al., 2017).

### Retrieval

To assess whether subsequent memory accuracy differed depending on the modality and perceived coherence of the narratives during encoding, we computed responses in each trial for every participant. Trials for which participants correctly indicated the picture depicting an event belonging to the cued narrative were labelled as “hit”. Trials in which participants did not choose the correct picture or did not provide an answer within the time limit were labelled as “miss”. Associated reaction times of correct trials were also computed for every participants. We fitted two separate linear mixed effects models using either correctness or reaction times in the retrieval task as the dependent variables. Modality (verbal and non-verbal) as well as coherence scores of each stimulus provided during encoding (from 1 to 5) were used as two fixed effect factors at the level of the individual trials. By-subject random intercept was further included in the two models to account for variability in the memory accuracy and reaction times at the level of individual participants. The model parameters were estimated using the restricted maximum likelihood approach.

### EEG statistical analysis

### Trial-based multiple regression analysis during narrative encoding

A trial-based multiple regression analysis was performed on time-frequency decomposition to assess how the position of the events predicted the power spectrum during the encoding of verbal and non-verbal narratives (**Figure 2**). To achieve this, the continuous power spectrum evoked during the 40-second narrative encoding was segmented based on the onset of each event composing the narrative., i.e., from the onset of the *first* event to the onset of the *second* event to create the “first event” epoch of the trial, then from the onset of the *second* event to the onset of the *third* event to create the “second event” epoch, and so on (**Figure 2C-1**). Because the duration of the events varied within the narratives and across stimuli, each power spectrum was fit with a b-spline interpolation approach to create an absolute time-frequency representation with a common matrix size of 30 Frequencies (from 1 to 30 Hz) and 126 time-points (from *n* event onset to event *n* offset, representing 100% of the absolute event length’s portion) at the original 500 Hz sampling resolution (**Figure 2C-2**). The power spectrum was then estimated using a general linear model including five regressors: (i) the modality of the narratives, (ii) the chronological position of each event within the narratives, (iii) the interaction between modality and position of events, (iv) the perceived coherence score of the narrative and (v) the Word2vec coherence scores as regressors. The first three regressors (i-iii) served to address our third hypothesis, while the perceived coherence and Word2vec coherence regressors (iv-v) reflected semantic parameters to account for participant- and event-related variance (**Figure 2C-3**). The beta weight of each regressor, obtained for every frequency x time point at each channel was standardised by dividing the standard error of the fit to obtain a t-value coefficient (Griffiths et al. 2021). Positive beta coefficients indicate that power spectrum increases with the values of a regressor (e.g., the chronological position of the events during the continuous encoding of the narratives), while negative values indicate a decrease with the values of a regressor. The spectral beta coefficients of each participant were then entered into a two-tailed cluster-based permutation test to assess whether every regressor predicted the observed power fits, by significantly deviating from the null hypothesis *t = 0* (2000 permutations, alpha threshold = 0.05, cluster alpha threshold = 0.05, minimum neighbourhood size = 3; probe values correction for two-tailed hypothesis; Maris and Oostenveld, 2007). The effect size of the significant clusters (*p*- value < 0.05) was quantified using Cohen’s *d_z_* (Lakens, 2013).

### Trial-based multiple regression analysis during narrative retrieval

The same trial-based multiple regression analysis was applied on the power spectrum when participants recollected narrative memories during the retrieval stage (“remember”). In this case, the power spectrum [1-30Hz] was modelled with a general linear model including the modality of encoding (verbal or non-verbal) as well as the coherence score of the narratives (1, 2, 3, 4 or 5) as regressors to predict it independently for every channel x frequency x time point. Similarly, the spectral beta coefficients of each participant were entered into two-tailed cluster-based permutation tests to assess whether the modality and the perceived coherence of the narratives during encoding predicted the observed power fits, by significantly deviating from the null hypothesis.

