## Supplementary Information for "Semantic elaboration determines the time course of alpha-beta oscillations during the encoding and retrieval of narrative memories"

### Behavioural performance during encoding and retrieval at single-trial level

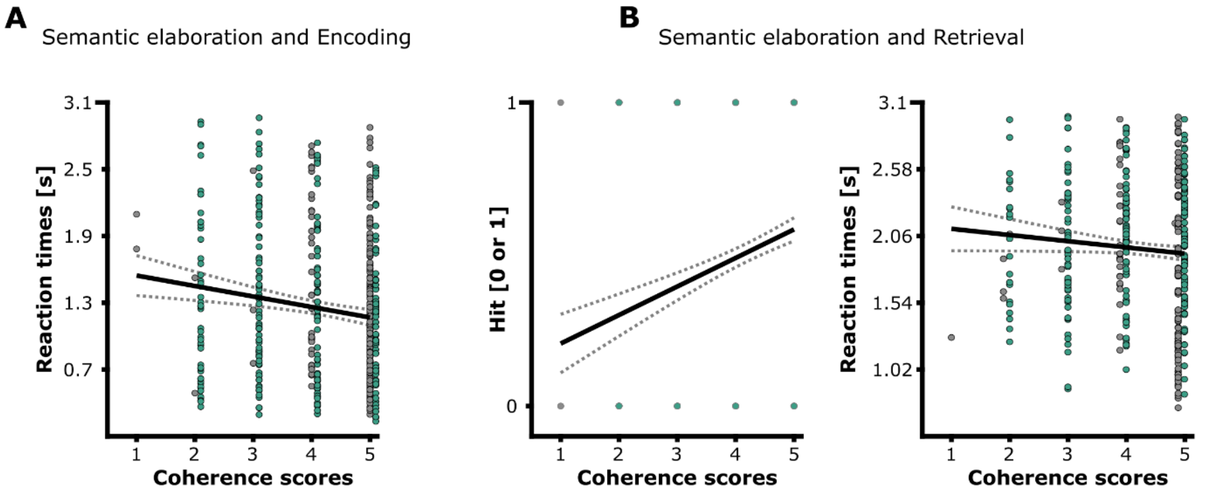


**Figure S1. Behavioural performance during encoding and retrieval.** (**A**) Effect of semantic elaboration on coherence perception speed during the encoding of verbal and non-verbal narratives. (**B**) Effect of semantic elaboration on subsequent memory accuracy and reaction times during verbal and non-verbal narrative retrieval. For each scatter plot, the solid line represents the linear regression model and the dashed curves show 95% confidence intervals for the fitted regression line. The dots represent one single trial for each stimulus across participants in the verbal (grey) and non-verbal (green) modalities.

### Positive relationship between perceived coherence scores of equivalent verbal and non-verbal narratives

We performed a linear regression model on the averaged coherence scores as well as reaction times across participants, between the narratives presented in verbal and equivalent non-verbal modalities (**Figure S2**). Results revealed a significant positive relationship for the perceived coherence scores across modalities, showing that verbal narratives with the highest scores were also highly rated in their non-verbal version ($F(1, 16) = 4.922$; $p = 0.041$; $R^2 = 0.188$). In contrast, no significant relationship was found for the reaction times ($F(1, 16) = 0.559$; $p = 0.465$; $R^2 = -0.027$).

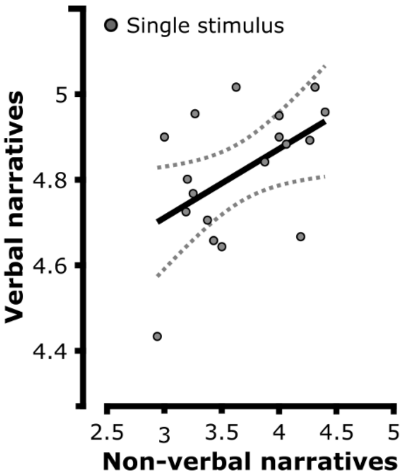


**Figure S2. Positive relationship between perceived coherence of corresponding verbal and non-verbal narratives.** Mean coherence scores of each narrative in the verbal (Y-axis) and equivalent non-verbal modality (X-axis) during encoding, averaged across participants.

### Word2vec coherence effect on power spectrum during narrative encoding

A cluster-based permutation was performed on the coefficients of the regressor Word2vec coherence score that reflected semantic parameters to account for event-related variance. Results revealed a significant positive cluster on power in the alpha-beta range, extending to lower theta range, and widespread over the scalp ($p$ = 0.002, cluster size = $9.12 \times 10^4$, mean t-statistic within cluster = 2.647; Cohen's $d_z$ = 0.483; **Figure S3**).

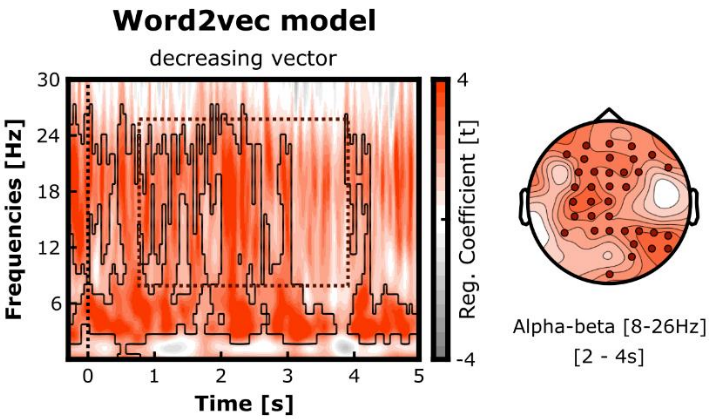


**Figure S3. Word2vec coherence effect on power spectrum during narrative encoding.** Time-frequency representation and topography of the effect of Word2vec coherence on power spectrum during narrative encoding (verbal and non-verbal trials collapsed together). Alpha-beta desynchronisation decreased with the semantic vector score of the events estimated by the Word2vec model (evidenced by the gradual darkness of the red colour in the plots). The time-frequency representations plot the t-values averaged across the significant channels (indicated by red dots on the topographies), and the black line outlines significant t-values. The topography shows the t-values for alpha-beta activity (8-26 Hz) within the time-window of interest outlined by the black line in the time-frequency plot.

#### Effect of narrative modality on alpha-beta oscillations during subsequent retrieval

Together with the perceived coherence scores, we also included the modality of encoding (verbal *vs.* non-verbal) as regressor to estimate the power spectrum during the retrieval with trial-based regression models.

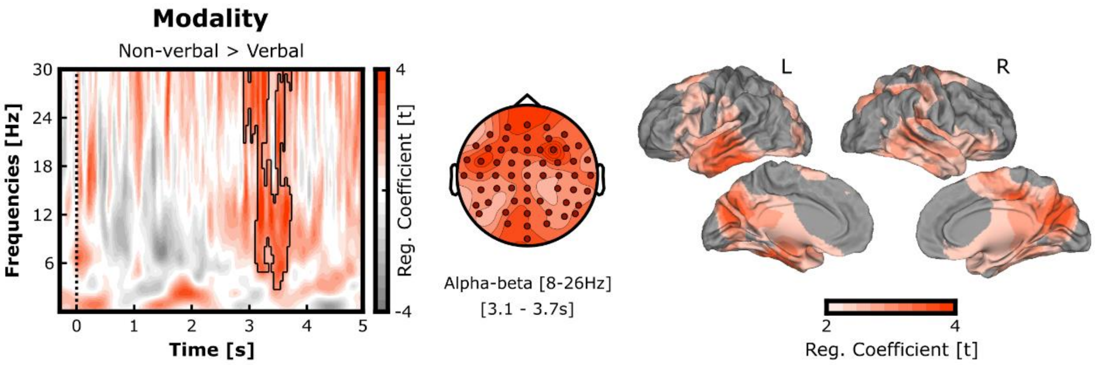


**Figure S4. Effect of narrative modality on alpha-beta oscillations during subsequent retrieval.** Left and centre: Time-frequency representation and topography of the effect of narrative modality on power spectrum during memory reinstatement (verbal vs. non-verbal). Alpha-beta oscillations desynchronized less during memory reinstatement when the narratives were encoded in the non-verbal compared to verbal modality (evidenced by the gradual darkness of the red colour in the plots). The time-frequency representation plots the t-values averaged across the significant channels, which are indicated by red dots on the topography. The black line outlines significant t-values. The topography shows the t-values for alpha-beta activity (8-26 Hz) within the time-window of interest [3.1 - 3.7s] outlined by the black line in the time-frequency plot. Right: Source localisation of the effect of narrative modality on alpha-beta oscillations in the time-window of interest (threshold at significant t-values).

Cluster-based permutation analysis performed on the regressor coefficients revealed a significant positive effect of modality of encoding on power in the alpha-beta range, extending to lower theta range, and widespread over the scalp ($p = 0.005$, cluster size = $1.294 \times 10^4$, mean t-statistic within cluster = 2.546; Cohen's $d_z = 0.465$; **Figure S4 left and centre**). Source analysis in the time-window of interest [3.1 - 3.7s] confirmed a positive correlation between narrative modality and alpha-beta power in the bilateral centro-parietal ($p = 4.996 \times 10^{-4}$, cluster size = $3.381 \times 10^3$, mean t-statistic within cluster = 2.713; Cohen's $d_z = 0.495$; **Figure S4 right**).

#### Alpha-beta power time-course during successful verbal and non-verbal memory retrieval

To complete our regressor model approach, we addressed the time-course of alpha-beta power when participants successfully retrieved memories of verbal and non-verbal narratives (**Figure S5**). Cluster-based permutation analysis performed on the difference of power spectrum revealed one early negative cluster ($p$ = 0.049, cluster size = -153.167, mean t-statistic within cluster = -2.553; Cohen's $d_z$ = 0.466) and one late positive cluster ($p$ = 0.023, cluster size = 201.027, mean t-statistic within cluster = 2.422; Cohen's $d_z$ = 0442). The difference of power between the two modalities was found in the centro-frontal regions, peaking in the expected alpha-beta frequency band, but also extending into the lower theta range of the spectrum [4-7Hz] (**Figure S5A**). The time-course of averaged alpha-beta power [8-26Hz] during the "remember" phase of the retrieval suggests that a similar desynchronisation followed by a resynchronisation rebound was driven by the successful recollection of verbal and non-verbal narrative memories, although delayed in the latter case. The corrected t-values for multiple comparisons confirmed two 'rebound' time windows occurring approximately from 2 to 3.5 seconds after the onset of "remember" (**Figure S5B**). Source localisation revealed a comparable early alpha-beta desynchronisation shortly occurring after the onset [0.2 - 0.8s] and localised in the bilateral occipital regions when participants retrieved either cued verbal ($p$ = 0.0085, cluster size = -1.372x$10^3$, mean t-statistic within cluster = -2.767; Cohen's $d_z$ = -0.505) or non-verbal narratives ($p$ = 0.008, cluster size = -1.2883x$10^3$, mean t-statistic within cluster = -3.188; Cohen's $d_z$ = -0.581**; Figure S5C**). The following alpha-beta rebound then occurred in the left temporal regions but with a different timing between verbal and non-verbal memories: permutation analysis revealed a significant positive cluster in the [1.5 - 2.1s] time-window for verbal ($p$ = 0.026, cluster size = 630.996, mean t-statistic within cluster = 2.32; Cohen's $d_z$ = 0.424) but not non-verbal retrieval. A second permutation analysis revealed a significant positive cluster later in the [2.8 - 3.4s] time-window for the non-verbal ($p$ = 0.017, cluster size = 921.535, mean t-statistic within cluster = 2.546; Cohen's $d_z$ = 0.465) but not the verbal retrieval.

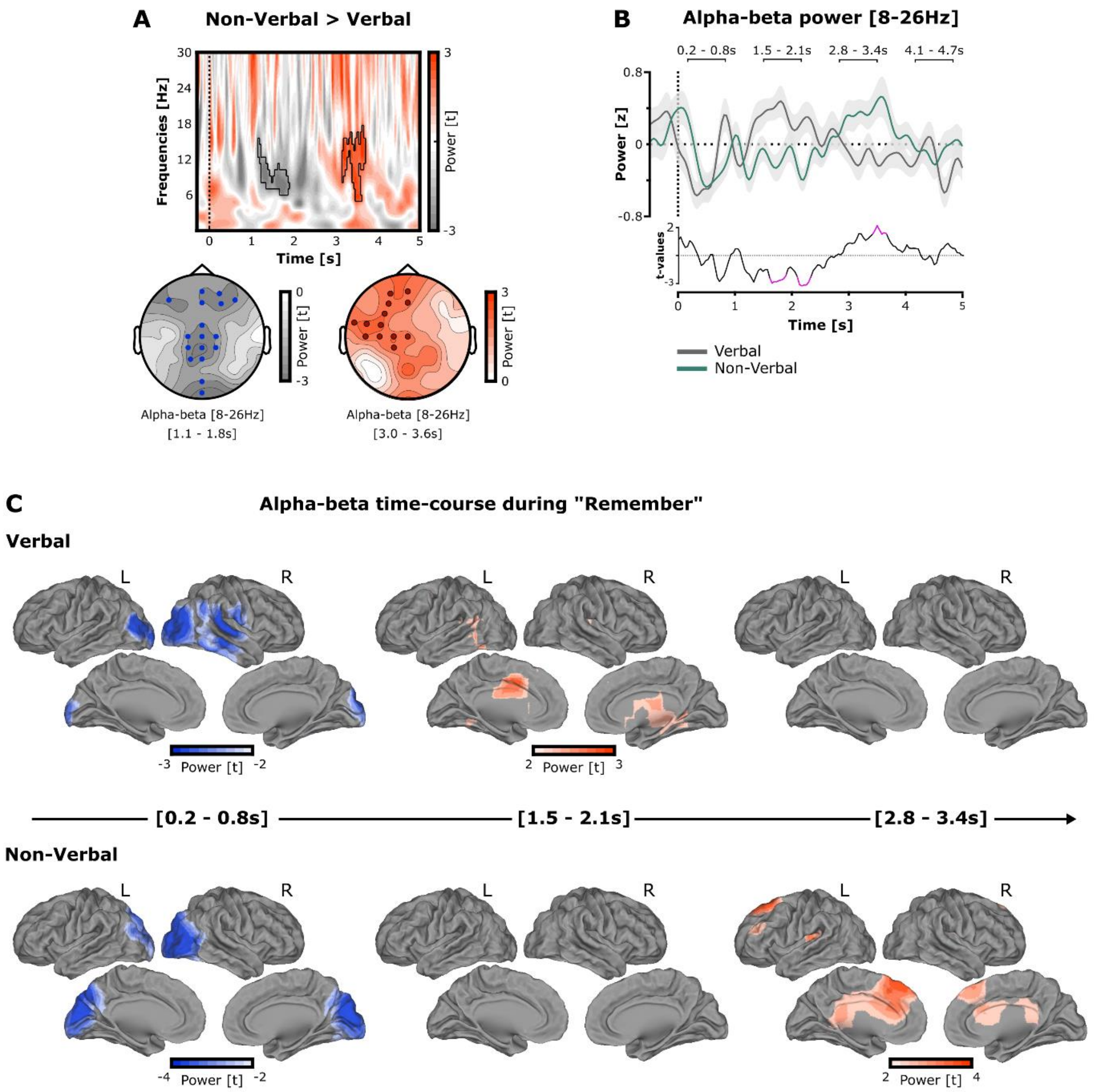


**Figure S5. Alpha-beta oscillation responses during the successful retrieval of verbal and non-verbal memories.** (**A**) Time-frequency representations and topographies of the power spectrum difference between verbal and non-verbal memories during the “remember” phase. A significant difference of alpha-beta power was found in the centro-frontal regions, being more synchronous first in the verbal then in the non-verbal modality during memory reinstatement. The time-frequency representation plots the t-values averaged across the significant channels, which are indicated by the blue and red dots on the topographies. The topographies show the t-values for alpha-beta activity (8-26 Hz) within the time-windows of interest [1.1 - 1.8s] and [3.0 - 3.6s] outlined by the black lines in the time-frequency plot. (**B**) The time-course of averaged alpha-beta power [8-26Hz] suggests that a similar desynchronisation followed by a “rebound” was driven by the successful reinstatement of narrative memories, although slower when participants successfully retrieved narratives encoded in the non-verbal modality (top). The t-values of alpha-beta power difference between the two modalities confirmed two 'rebound' time-windows occurring approximately from 2 to 3.5 seconds after the onset of “remember” (bottom; significant t-values corrected for multiple comparisons plotted in pink colour). (**C**) Source localisation of the alpha-beta power modulation during verbal (top) and non-verbal (bottom) narrative memory retrieval (threshold at significant t-values). The alpha-beta activity for memory reinstatement was split in three time-windows of interest driven by the statistical differences from panel B: early desynchronisation [0.2 - 0.8s], rebound in the verbal modality [1.5 - 2.1s] and rebound in the non-verbal modality [2.8 - 3.4s].
